## Supplementary figures for "Domain-adaptive neural networks improve supervised machine learning based on simulated population genetic data"

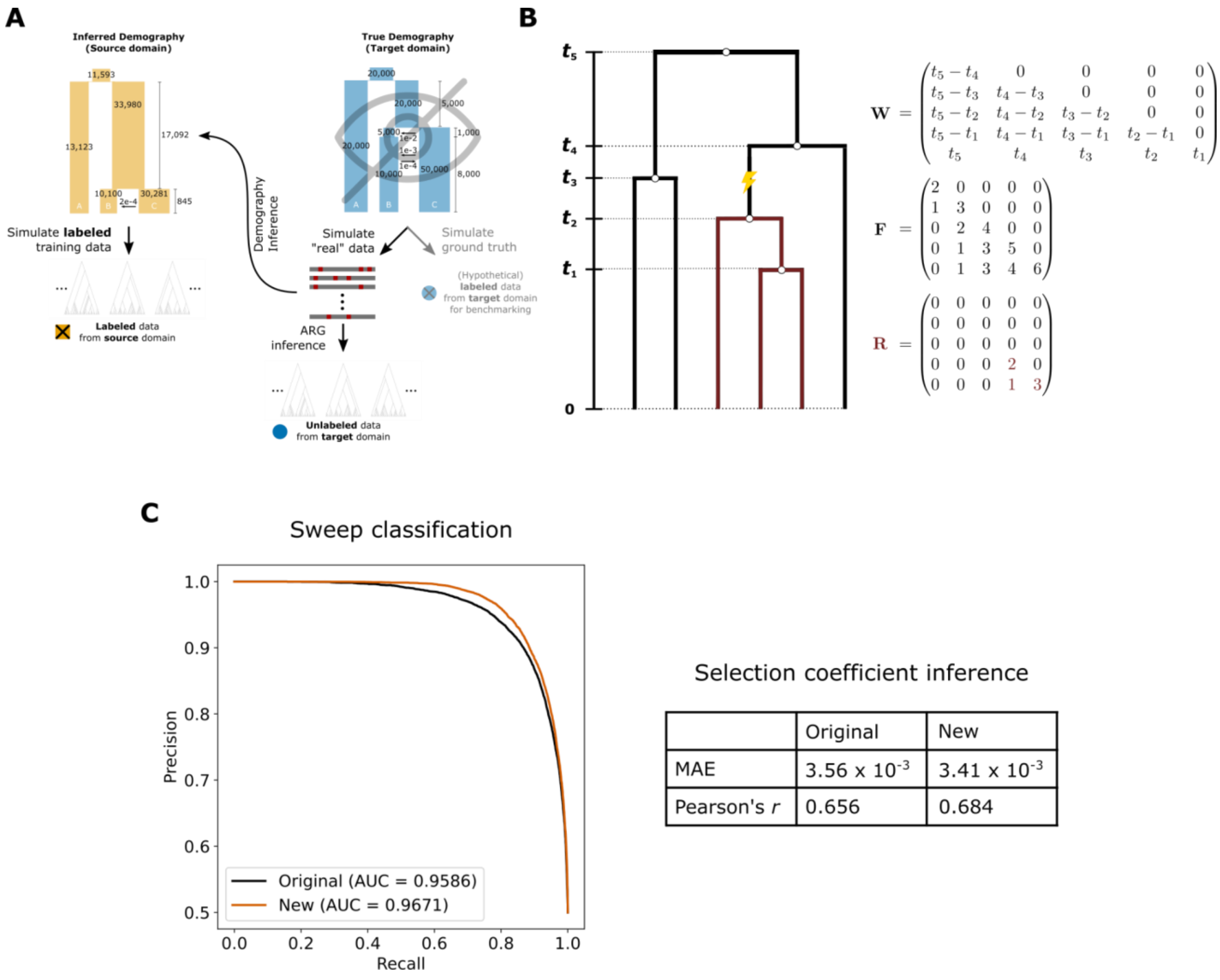


Selection coefficient inference

| | Original | New |
|---|---|---|
| MAE | $3.56 \times 10^{-3}$ | $3.41 \times 10^{-3}$ |
| Pearson's *r* | 0.656 | 0.684 |

**Supplementary Figure 1. Domain-adaptive SIA. A)** The workflow of a simulation study that aims to benchmark the performance of the domain-adaptive SIA model in a realistic setting of demographic mis-specification. **B)** An improved version of SIA input features that encodes the full genealogy (adapted from [59]). A genealogy with *n* taxa at a polymorphic site is uniquely encoded by three (*n*-1) x (*n*-1) lower triangular matrices. The weight matrix **W** encodes the coalescent intervals where $w_{ij} = t_{n-j} - t_{n-1-i}, \forall\, i \geq j$, and the topology matrix **F** encodes the number of lineages persistent in the coalescent intervals *corresponding* to **W** (i.e. $f_{ij} = \#\, of\ lineages\ between\ t_{n-j} and\ t_{n-1-i}, \forall i \geq j$). The derived lineage matrix **R** encodes only the subtree subtending the branch where the mutation occurred (red lightning symbol), following the same scheme as **F**. Note that the **W** matrix is a redundant encoding of the *n*-1 coalescent times ($t_1$,$t_2$, …,$t_{n-1}$), which contains information roughly equivalent to the original SIA input features [12]. **C)** Comparison of the performance of the new SIA input features in (**B**) to that of the original SIA input features.

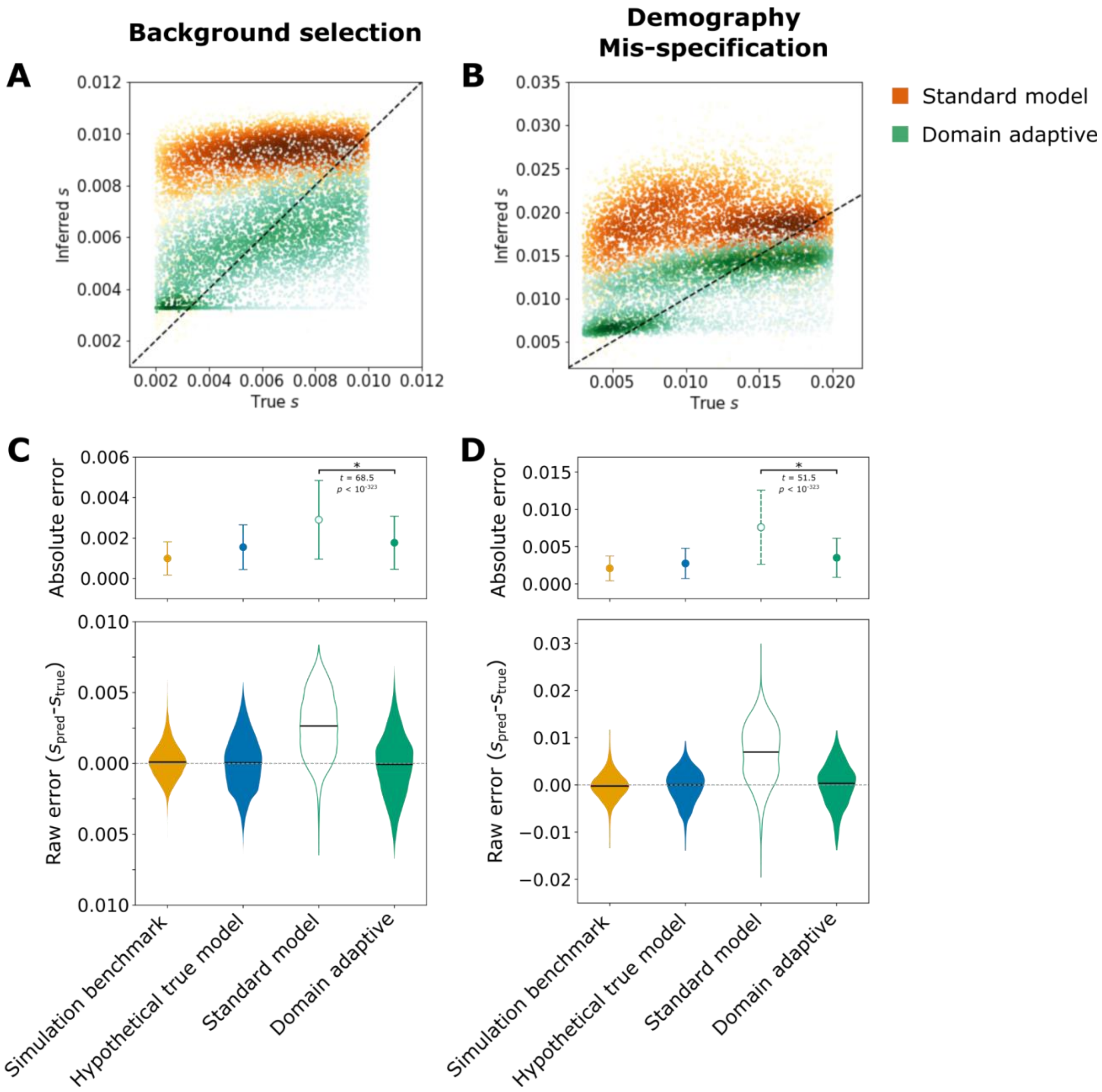


**Supplementary Figure 2. Selection coefficient inference performance of SIA models.** Raw data used to plot **Figs. 3B** and **3D** are presented in (**A**) and (**B**), respectively. Performance of SIA models in the simulation experiment of failure to account for background selection (**C**) and in the simulation experiment of demographic model mis-specification (**D**) is presented in terms of mean and standard deviation of the absolute error (top) as well as the distribution of raw error (bottom). Statistical significance (*) of the difference between the absolute error of the standard model and that of the domain-adaptive model is evaluated with Welch's *t*-test. See **Fig. 1C** for definition of the model labels.

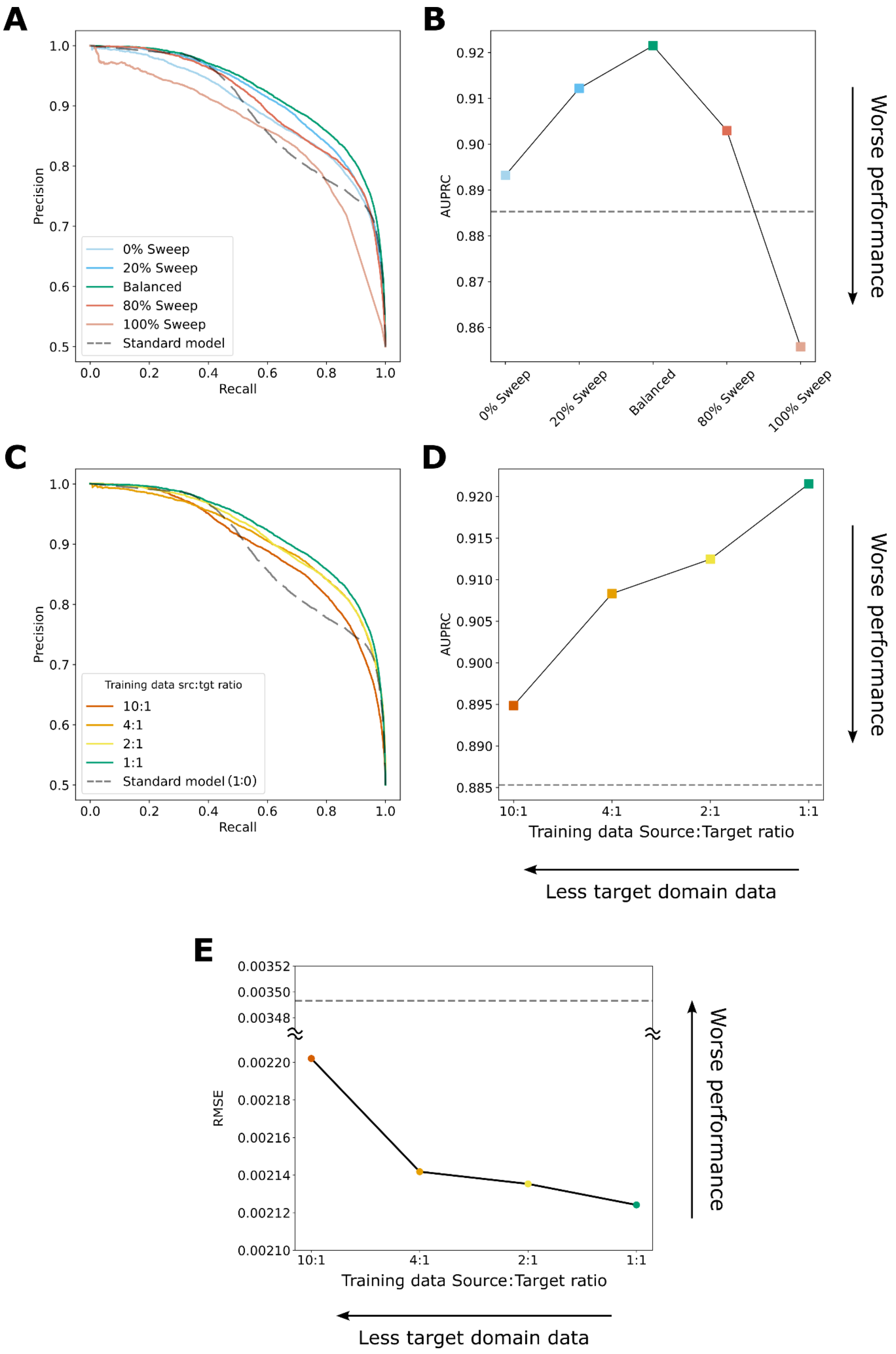

A
B
C
D
E
Precision
Recall
0% Sweep
20% Sweep
Balanced
80% Sweep
100% Sweep
Standard model
AUPRC
Worse performance
Training data src:tgt ratio
10:1
4:1
2:1
1:1
Standard model (1:0)
Training data Source:Target ratio
Less target domain data
RMSE

**Supplementary Figure 3. Performance of dadaSIA models trained with imbalanced data.** The sweep classification performance of dadaSIA models trained with different proportions of sweep vs. neutral examples in the target domain is shown in the form of precision-recall curves (**A**) and the area under precision-recall curve (AUPRC) (**B**). Note that the performance is always evaluated on a balanced test set. The performance of dadaSIA models trained with less target domain data than source domain data is shown in the form of precision-recall curves (**C**) and the values of AUPRC (**D**) for the classification task, and in the form of root mean squared error (RMSE) (**E**) for the selection coefficient inference task. The dashed lines in (**B**), (**D**) and (**E**) indicate performance of the standard model.

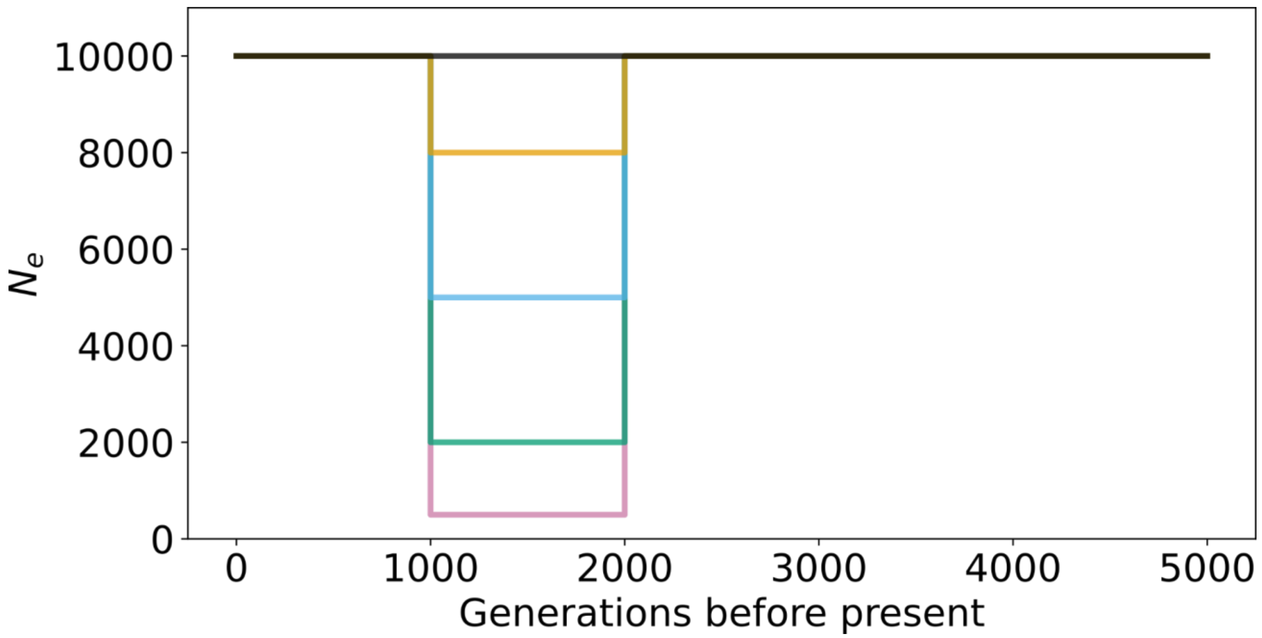


**Supplementary Figure 4. Demographic mis-specification in the form of different degrees of bottlenecks tested in Fig. 5 experiments.**

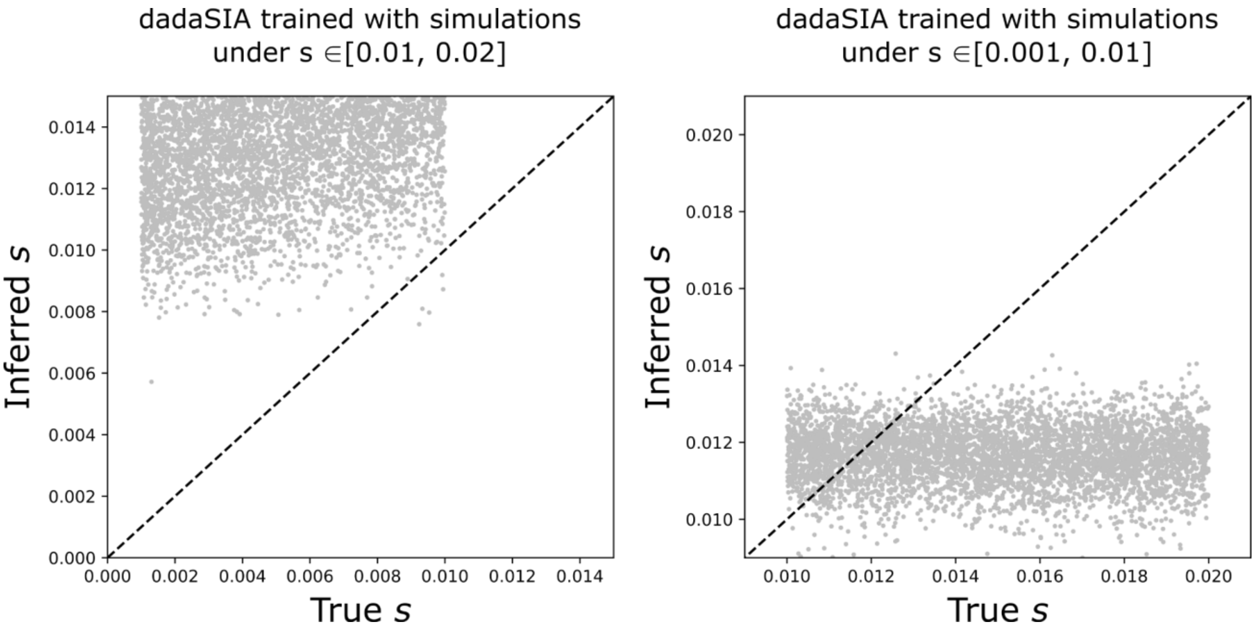


**Supplementary Figure 5**. **Inference of out-of-range selection coefficients in the target domain using the dadaSIA model.** The dadaSIA model trained with source domain data under $s \in [0.01, 0.02]$ failed to meaningfully infer any value lower than 0.01, even when examples of $s \in [0.001, 0.01]$ were supplied to the model as “unlabeled” target domain data, and vice versa.

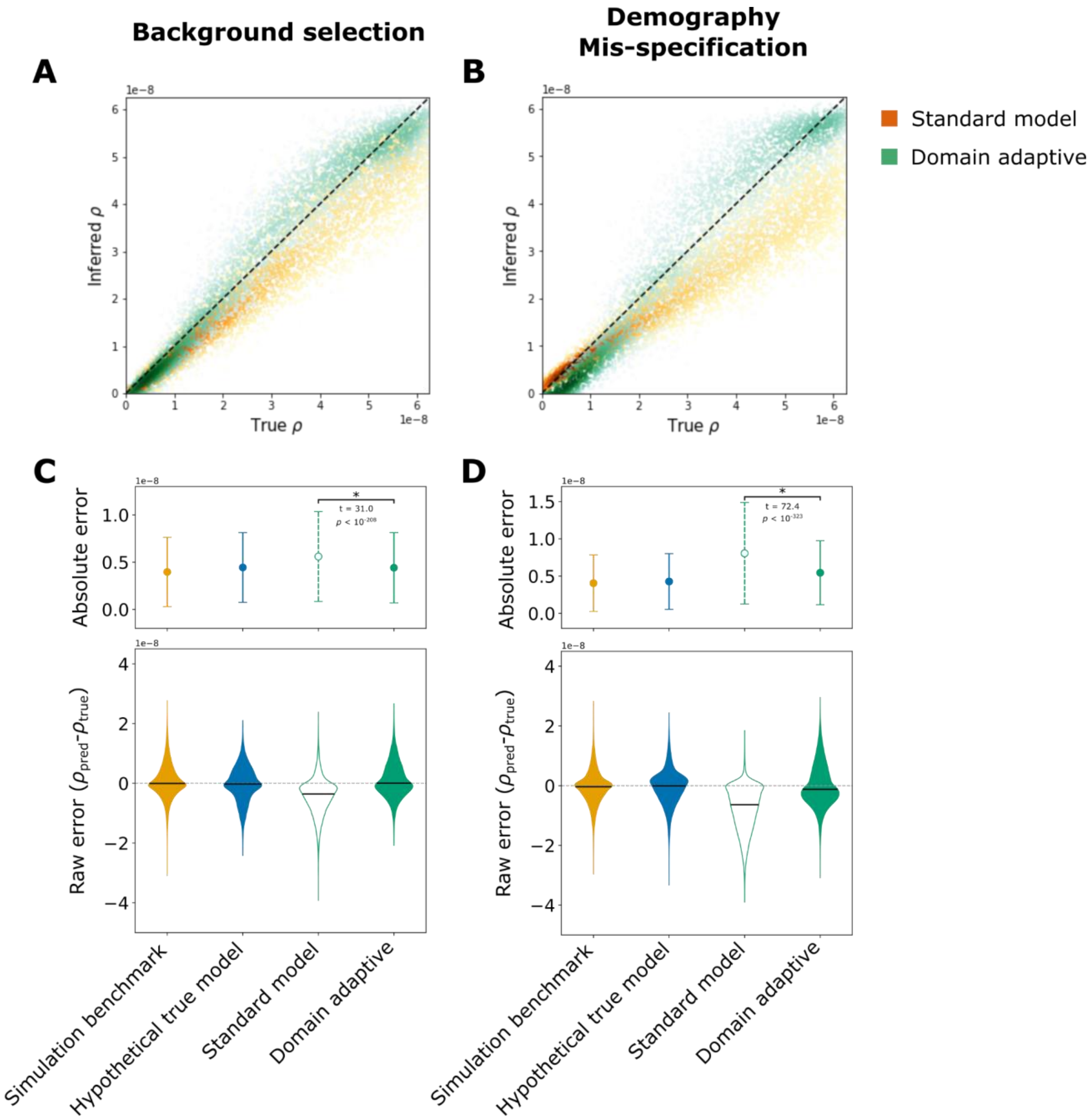


**Supplementary Figure 6**. **Recombination rate inference performance of ReLERNN models.** Raw data used to plot **Figs. 4A** and **4B** are presented in (**A**) and (**B**), respectively. Performance of ReLERNN models in the simulation experiment of failure to account for background selection (**C**) and in the simulation experiment of demographic model mis-specification (**D**) is presented in terms of mean and standard deviation of the absolute error (top) as well as the distribution of raw error (bottom). Statistical significance (*) of the difference between the absolute error of the standard model and that of the domain-adaptive model is evaluated with Welch's *t*-test. See **Fig. 1C** for definition of the model labels.

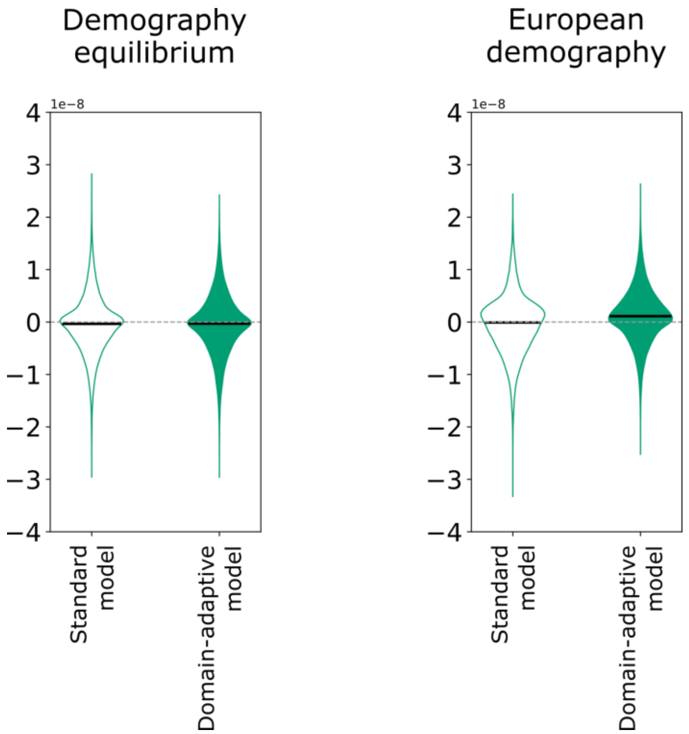


**Supplementary Figure 7. Distribution of raw error of the ReLERNN models inferring recombination rate without simulation mis-specification.** The respective mean absolute error (MAE) of the standard and domain-adaptive models are 4.05 x $10^{-9}$ and 4.13 x $10^{-9}$, under demography equilibrium, and 4.28 x $10^{-9}$ and 3.93 x $10^{-9}$, under a European demography. Note that the domain-adaptive model has a slight upward bias in its estimates in the case of European demography.

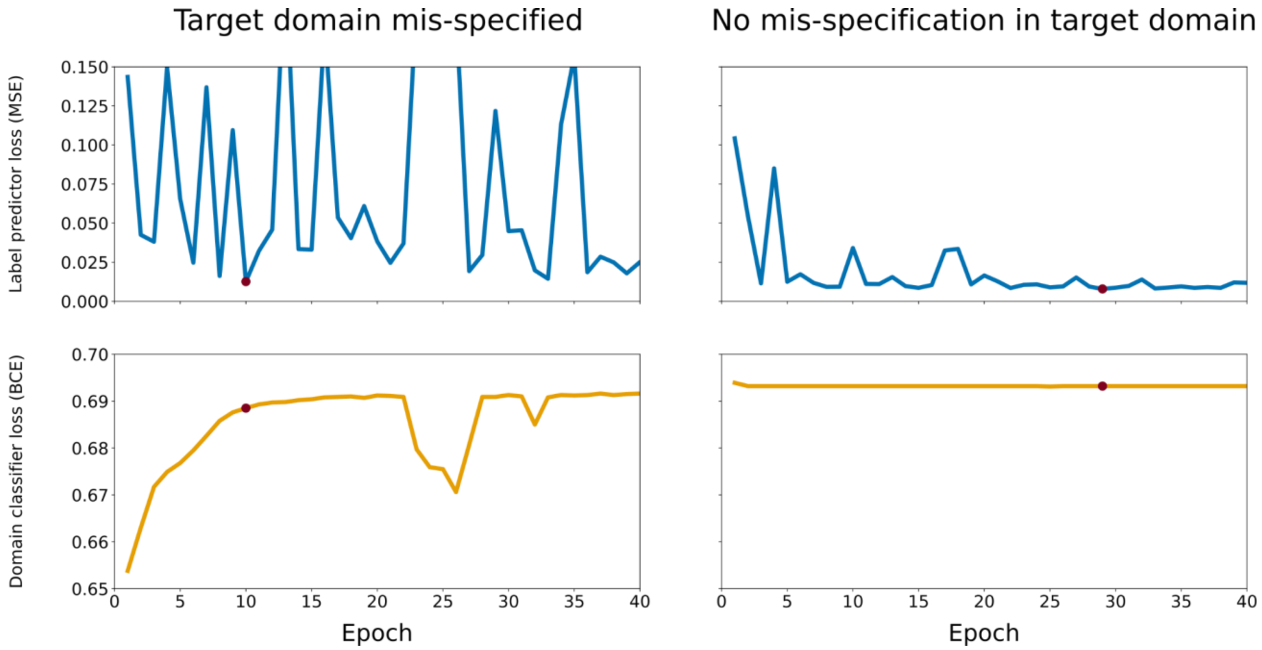


**Supplementary Figure 8. Validation loss of the label predictor branch (mean squared error) and the domain classifier branch (binary cross entropy) over training epochs.** The losses of the domain-adaptive ReLERNN models during training are plotted with and without simulation mis-specification. The red dot marks the early-stopping epoch (i.e. epoch with the lowest validation loss for the label predictor).

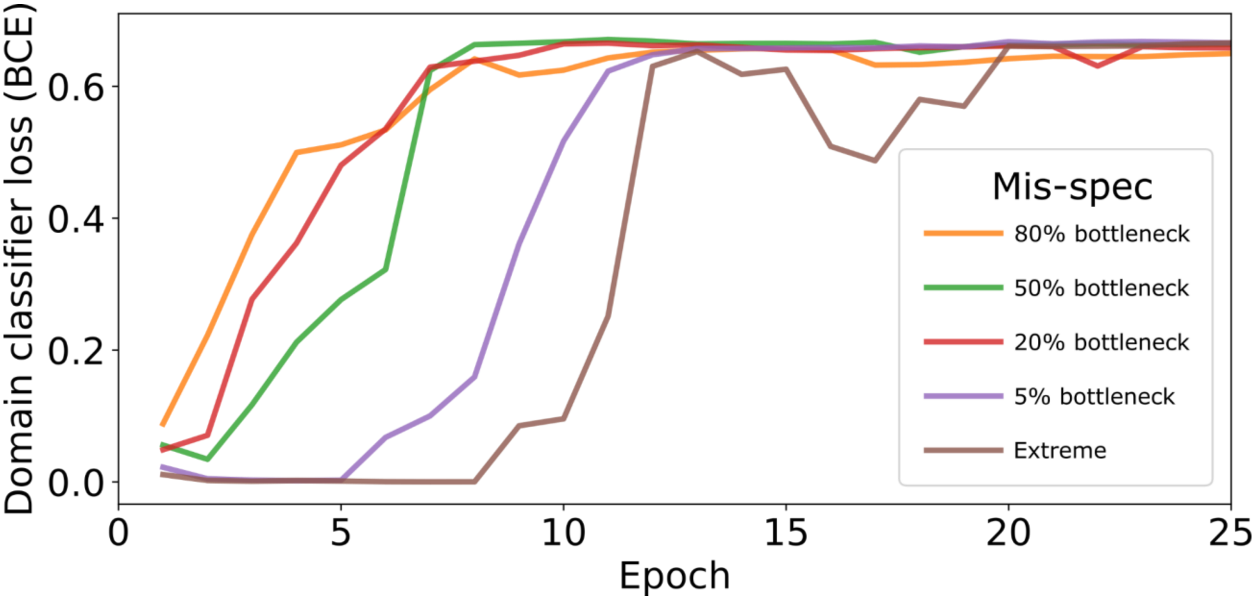


**Supplementary Figure 9**. **Domain classifier loss of dadaSIA models under different degrees of simulation mis-specification.** See **Fig. 5** and **Methods** for details of the types of mis-specification.
